# Cleavage of a pathogen apoplastic protein by plant subtilases activates immunity

**DOI:** 10.1101/2019.12.16.878272

**Authors:** Shuaishuai Wang, Rongkang Xing, Yan Wang, Haidong Shu, Shenggui Fu, Judith K. Paulus, Mariana Schuster, Diane G.O. Saunders, Joe Win, Vivianne Vleeshouwers, Xiaobo Zheng, Renier A. L. van der Hoorn, Sophien Kamoun, Suomeng Dong

**Affiliations:** Department of Plant Pathology, Nanjing Agricultural University, Nanjing, China; The Plant Chemetics Laboratory, Department of Plant Sciences, University of Oxford, South Parks Road, Oxford OX1 3RB, United Kingdom; The Sainsbury Laboratory, University of East Anglia, Norwich Research Park, Norwich NR4 7UH, United Kingdom; John Innes Centre, Norwich Research Park, Norwich, NR4 7UH, United Kingdom; Wageningen UR Plant Breeding, Wageningen University and Research Centre, Droevendaalsesteeg 1, Wageningen 6708 PB, The Netherlands

## Abstract

The plant apoplast is a harsh environment in which hydrolytic enzymes, especially proteases, accumulate during pathogen infection. However, the defense functions of most apoplastic proteases remains largely elusive. Here, we show that a newly identified small cysteine-rich secreted protein PC2 from the potato late blight pathogen *Phytophthora infestans* induces immunity in Solanum plant species only after cleavage by plant apoplastic subtilisin-like proteases, such as tomato P69B. A minimal 61-amino-acid core peptide carrying two key cysteines and widely conserved among most oomycete species is sufficient for PC2 activity. Kazal-like protease inhibitors, such as EPI1 produced by *P. infestans* can prevent PC2 cleavage and dampen PC2 elicited host immunity. This study reveals that cleavage of pathogen proteins to release immunogenic peptides is an important function of apoplastic proteases but that pathogens interfere with these functions using protease inhibitor effectors.

## Introduction

The plant apoplast is the extracellular compartment serving as a major battlefield for plant-pathogen interactions(Du et al., 2016). To defend against invading microbes, plants have evolved pattern recognition receptors (PRRs) to recognize microbe-associated molecular patterns (MAMPs) in the apoplastic interface and activate innate immunity(Boller and He, 2009; Miao et al., 2019). MAMPs comprise a wide category of molecules such as bacterial flagellin, elongation factors, peptidoglycan, and fungal cell wall chitin(Newman et al., 2013). Recently, apoplastic proteins have been recognized as an exciting resource for identifying new MAMPs. Some of these proteins are effectors that act to break down plant extracellular immunity through their biochemical functions. However, the peptide signatures of these proteins can also be recognized by plant membrane-localized PRRs to trigger a series of immune responses including reactive oxygen species (ROS) production, defense-related marker gene expression or hypersensitive cell death(Jones and Dangl, 2006). These proteins such as necrosis and ethylene-inducing protein (NEP) and xyloglucan-specific endoglucanase (XEG) are evolutionarily conserved across microbial taxa, providing an opportunity to study broad spectrum plant-microbe apoplastic interactions(Gijzen and Nurnberger, 2006; Ma et al., 2015). However, the mode of action of most apoplastic proteins remains largely elusive. Revealing the mechanisms at play in the plant apoplastic space during microbial infection can provide novel insight into pathogenesis and identify new targets for engineering more robust plant immunity.

The plant apoplast is a harsh environment which is full of digestive enzymes including proteases. A few plant proteases are highly up-regulated during pathogen infection and have long been tied to plant immunity. One such example is the tomato subtilisin-like serine protease P69B, which was first reported as a pathogenesis-related (PR) protein(Tornero et al., 1997). Other evidence pinpoints apoplastic proteases as important players in plant immunity. For instance, silencing the papain-like cysteine protease (PLCP) C14 orthologs in Nicotiana benthamiana leads to increased plant susceptibility to *Phytophthora infestans*, the causal agent of potato late blight(Kaschani et al., 2010). Moreover, depletion of tomato PLCP PIP1 results in hyper-susceptibility to fungal, bacterial, and oomycete diseases, demonstrating that PIP1 constitutes a basal defense to a wide range of pathogens(Ilyas et al., 2015). Apoplastic proteases act in plant immunity in either protease activity-dependent or independent manners. Rcr3, another tomato PLCP, is essential for the recognition of the receptor Cf-2 to the apoplastic effector Avr2 from *Cladosporium fulvum*, which acts as a PLCP inhibitor(Krüger et al., 2002). However, inhibition of Rcr3 activity using the PLCP inhibitor E-64 does not trigger Cf-2–mediated HR, suggesting Cf-2 triggered plant immunity is independent of Rcr3 activity(Rooney et al., 2005). In contrast, a recent study demonstrates that the release of the maize immune signaling peptide Zip1 requires maize PLCPs. In this case, PLCP activity activates SA immune signaling and maize resistance to the maize pathogen *Ustilago maydis*(Ziemann et al., 2018). Plant genomes encode a large family of proteases with distinct predicted activities. Identifying the substrates of these proteases is critical to uncover the mode of action of these proteases. However, the substrates of most proteases remain unclear. Although the underlying mechanisms are far from clear, current findings demonstrate that plant proteases play an important role in the regulation of plant immunity.

Inhibition of plant proteases represents a general counter-defense strategy used by invading pathogens(Figueiredo et al., 2018). To achieve successful colonization, adapted microbial pathogens deliver protease inhibitors into the host apoplast to counteract host defense. For example, fungal apoplastic effector Pit2 secreted by U. maydis inhibits a set of apoplastic maize cysteine proteases implicated in maize defense and contributes to the virulence of *U. maydis* in maize(Mueller et al., 2013; Misas Villamil et al., 2019). In Phytophthora, a well-characterized group of intercellular proteins are the extracellular cystatin-like protease inhibitors (EPIC). *P. infestans* EPIC1 binds to tomato PLCP Rcr3 and C14 and inhibits their activity in the tomato apoplast(Kaschani et al., 2010). EPIC2B is a more robust cysteine protease inhibitor with stronger inhibitory activity against Rcr3 and Pip1(Dong et al., 2014). Beside Phytophthora EPIC2B and fungal Avr2, Gr-Vap1 from cyst nematode and Cip1 from Pseudomonas bacteria pathogen also target these tomato proteases(Misas Villamil et al., 2019), indicating that different pathogens have independently evolved divergent effectors to inhibit plant protease during co-evolution. In addition, *P. infestans* secretes Kazal-like serine protease inhibitors, including the EPI1 and EPI10 proteins that inhibit tomato subtilase P69B(Tian et al., 2004; Tian et al., 2005). More recently, a study on the pathogenicity of Huanglongbing uncovered the Candidatus Liberibacter asiaticus effector SDE1 that directly interacts with and inhibits citrus PLCP activity(Clark et al., 2018). In another case, the cyst nematode Heterodera schachtii effector protein 4E02 repurposes a PLCP from its role in defense by targeting it to distinct plant cell compartments without inhibibiting its activity(Pogorelko et al., 2019). These studies illustrate a concerted interaction between the host proteases and the pathogen protease inhibitors, which dictates the success of pathogen colonization.

Interestingly, one common signature of apoplastic proteins is the presence of cysteine residues, which likely form disulfide bonds within the effectors to maintain their stability and biochemical function under harsh plant extracellular conditions. Early study on fungal avirulence effectors led to the characterization of small cysteine-rich (SCR) proteins which are typified by their rich content of cysteine residues and small molecular weight(Templeton et al., 1994). In other plant pathogens such as oomycetes, SCR proteins form a major group of apoplastic effectors(Torto et al., 2003). PcF, an SCR effector secreted by *Phytophthora cactorum* induces necrosis in plants(Orsomando et al., 2001). Another SCR effector SCR74 was identified in *P. infestans*. Although the function of SCR74 remains unclear, the genes encoding SCR74 family of effectors are under strong diversifying selection(Liu et al., 2005).

In this study, we developed a pipeline to identify potential SCR proteins in *P. infestans* and performed a high-throughput SCR proteins functional screen in N. benthamiana for immune response. We identified the *P. infestans* SCR protein PC2, that is capable of triggering plant immune responses in a manner typicle of MAMP. Further investigation revealed that PC2 is proteolytically cleaved by plant serine proteases such as tomato P69B. Genetic manipulation of P69B and chemical inhibition of serine protease activity both act to impair PC2-triggered immunity, suggesting that the cleavage process is essential for PC2-triggered plant immunity. In addition, we found that the cleavage could be suppressed by EPIs, the *Phytophthora* Kazal-like protease inhibitors. This study highlights an example of defense-counterdefense mechanisms in plant-microbe interactions with a finding that a plant protease cleaves a pathogen apoplastic protein to activate defense responses to achieve immunity, and the pathogen deploys protease inhibitors as a counter-defense measure to evade recognition by the host.

## Results

### Identification of the *P. infestans* SCR protein PC2

To identify candidate *P. infestans* apoplastic proteins as MAMPs for further investigation, we established a bioinformatic pipeline to predict putative apoplastic proteins from the *P. infestans* genome(Saunders et al., 2012). A total of 65 SCR proteins were predicted based on characteristics typical of apoplastic effectors such as the presence of a secretion signal, a high number of cysteine residues, and small protein size (Figure 1A, Supplementary Table 1). Among these genes, 37 encode known elicitin, CBEL, and protease inhibitor homologs. Next, we evaluated the expression of the other 28 candidate SCR proteins in publicly available gene expression data(Gajendran et al., 2006). This analysis identified a subset of 5 genes encoding predicted SCR protein that were highly expressed during mycelium, sporangia, and infection stages (Supplementary Table 2). Therefore, we focused on the remaining 5 candidate SCR proteins for further evaluation.

**Figure 1.**
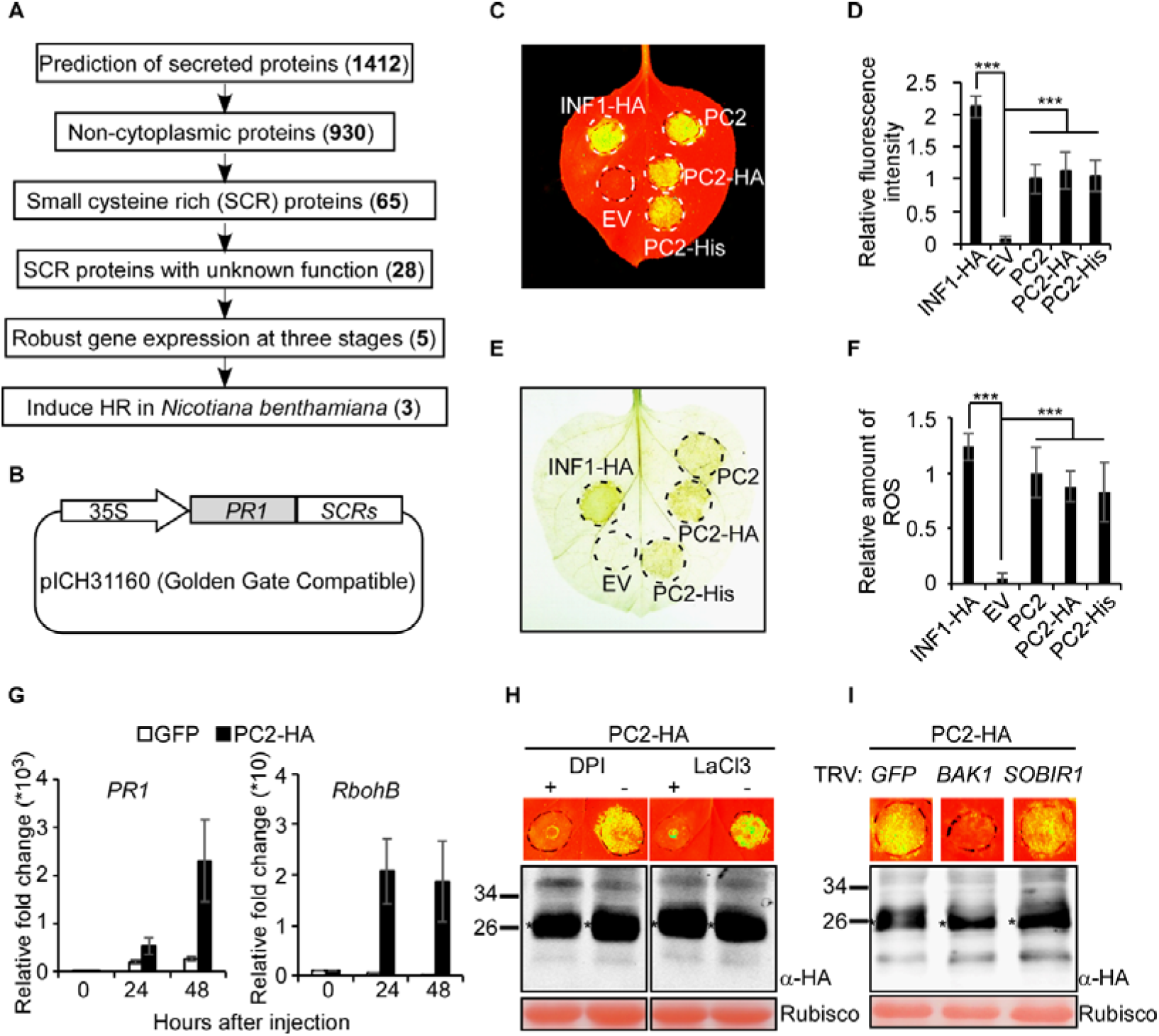
Screening of *P. infestans* SCR proteins led to the discovery of PC2. (**A**) Screening pipeline for small cysteine-rich proteins from *P. infestans*. (**B**) Schematic drawing of the plant expression constructs for candidate SCR effectors. The N-terminal signal peptide of all these effectors was deleted and replaced by the PR1 signal peptide. (**C**) Representative *N. benthamiana* leaves 5 d post inoculation with agrobacterium strains expressing PC2, PC2-His or PC2-HA cloned in PVX vector, along with the empty vector (EV). INF1-HA was used as a positive control. (**D**) Quantitative fluorescence intensity of cell death. Means from five independent replicates are shown (***P< 0.001; one-way ANOVA). (**E**) ROS accumulation at 2.5 dpi in *N. benthamiana* leaves infiltrated with agrobacteria containing PC2, PC2-His, PC2-HA or EV, as detected by DAB staining. (**F**) Quantitative intensity of DAB staining. Means and standard errors from five independent replicates (***P< 0.001; one-way ANOVA). (**G**) Transcript level of *PR1* and *RbohB* induced by PC2 in *N. benthamiana* leaves. The gene expression was examined at different time points by using qRT-PCR. Means and standard errors from three independent replicates are shown. (**H**) PC2-induced cell death in *N. benthamiana* leaves was inhibited by DPI and Lacl_3_. (**I**) BAK1 is required for PC2-triggered cell death in *N. benthamiana*. *N. benthamiana* plants were subjected to VIGS by inoculation with TRV constructs (TRV:*GFP*, TRV:*BAK1*, and TRV:*SOBIR1*). Four weeks after inoculation, PC2 was transiently expressed in the gene-silenced leaves and then leaves were photographed 5 days later. The experiment was performed three times with three plants for each TRV construct.

To investigate the biological function of these 5 candidate SCR proteins in planta, we carried out the ectopic expression assay in *N. benthamiana*. First, each gene was synthesized without its native secretion signal and codon optimized for in planta expression. All sequences were assembled with the 35S promoter and *Nicotiana* pathogenesis-related protein 1 (PR1) signal peptide for secretion (Figure 1B). Effector expression in *N. benthamiana* utilized the potato virus X (PVX) transient expression system mediated by *Agrobacterium tumefaciens*. At five days post-infiltration, ectopic expression of 3 candidate SCR genes triggered cell death in *N. benthamiana* (Supplementary Table 2). Among these genes, PITG_14439 (designated PC2) encodes an SCR protein containing 18 cysteine residues and has homologs among all the *Phytophthora* species we examined so far. To further evaluate the expression of PC2 during infection we carried out qRT-PCR. *P. infestans* strain T30-4 was used to inoculate Heinz tomato leaves and samples were collected at nine time points ranging from 3 hpi to 120 hpi. Subsequent qRT-PCR analysis illustrated that PC2 is up-regulated during infection, peaking at 48 hours post infection (Supplementary Figure 1A). The *Phytopthora* marker gene Avr3a and NPP1.1 clearly show early and late expression pattern respectively in our assay (Supplementary Figure 1B).

To study the biological activity of PC2 protein, we transiently expressed PC2 with HA or His tags and evaluated its role in inducing cell death in various Solanum species. The well-studied elicitor gene *INF1* and necrosis gene *NLP1* served as positive controls. Transient expression of PC2 in *N. benthamiana*, *N. tabacum*, *Solanum lycopersicum*, *S. melongena*, and *S. tuberosum* induced typical cell death or necrosis symptoms (Figures 1C and 1D; Supplementary Figure 2). These results suggest that PC2 is able to trigger cell death in a wide range of Solanum species. To further evaluate PC2-triggered plant responses, we examined ROS accumulation and the expression of *PR* genes after PC2 expression in *N. benthamiana*. Staining with 3,3’-diaminobenzidine (DAB) at 2.5 dpi showed that PC2 expression leads to a high level of ROS accumulation (Figures 1E and 1F). In addition, RT-qPCR assays demonstrated that the *N. benthamiana* defense-related marker genes *NbPR1* and *NbRbohB* were significantly up-regulated in response to PC2 expression (Figure 1G). Furthermore, the application of the ROS inhibitor DPI or calcium channel inhibitor Lacl_3_ blocked PC2-induced cell death (Figure 1H).

Next we evaluated whether PC2-induced immunity requires a PRR co-receptor by evaluating PC2-induced cell death in *BAK1* and *SOBIR1* silenced *N. benthamiana* plants(Heese et al., 2007; Liebrand et al., 2013) (Supplementary Figure 3). PC2-induced cell death was only impaired in *BAK1-* but not *SOBIR1-* silenced plants (Figure 1I), indicating that *N. benthamiana* perception of PC2 likely requires the PRR co-receptor BAK1. Together these results indicate that PC2-induced immune responses require calcium burst and ROS production and highly dependent on plant membrane-associated LRR-RK as PRR.

To further examine the role of PC2 during infection, we overexpressed *PC2* in *P. infestans* (Supplementary Figure 4A) and evaluated three independent transformants (T8, T14, and T29). All transformants showed normal filamentous growth when compared to the WT recipient strain (Supplementary Figure 4B). Expression analysis by RT-qPCR demonstrated that *PC2* expression was elevated in each of the three transformants (Supplementary Figure 4C). To measure their virulence, mycelia from each transformant and the recipient strain were inoculated onto leaves of *S. tuberosum* cultivar Desiree (Supplementary Figures 4D and 4E) and *N. benthamiana* (Supplementary Figures 4F and 4G). All three *P. infestans* transformants produced significantly smaller infection lesions compared to those of the wild-type recipient strain, indicating that misexpression of PC2 impairs *P. infestans* infection.

### *PC2* orthologs are widely present in oomycete species

To determine the presence of PC2 orthologs in oomycetes, we performed a blastp search in an oomycete proteome database (Supplementary Table 3). *PC2* orthologous genes were identified in *Phytophthora*, downy mildew, *Salisapilias*, and *Pythium* genomes. The proteins encoded by these homologous genes varied in length, but all carried signal peptides and conserved cysteine residues (Figure 2A, Supplementary Figure 5). However, we failed to detect any homologous sequence from *Albugo* plant pathogen genomes. Phylogenetic analysis revealed that PC2 and its orthologs from *Phytophthora* spp. and *Peronophythora* spp. form a clade distinct from those of other oomycetes, with the orthologs from *H*. *arabidopsidis* and *Pythium* being the least related relatives (Figure 2A).

**Figure 2.**
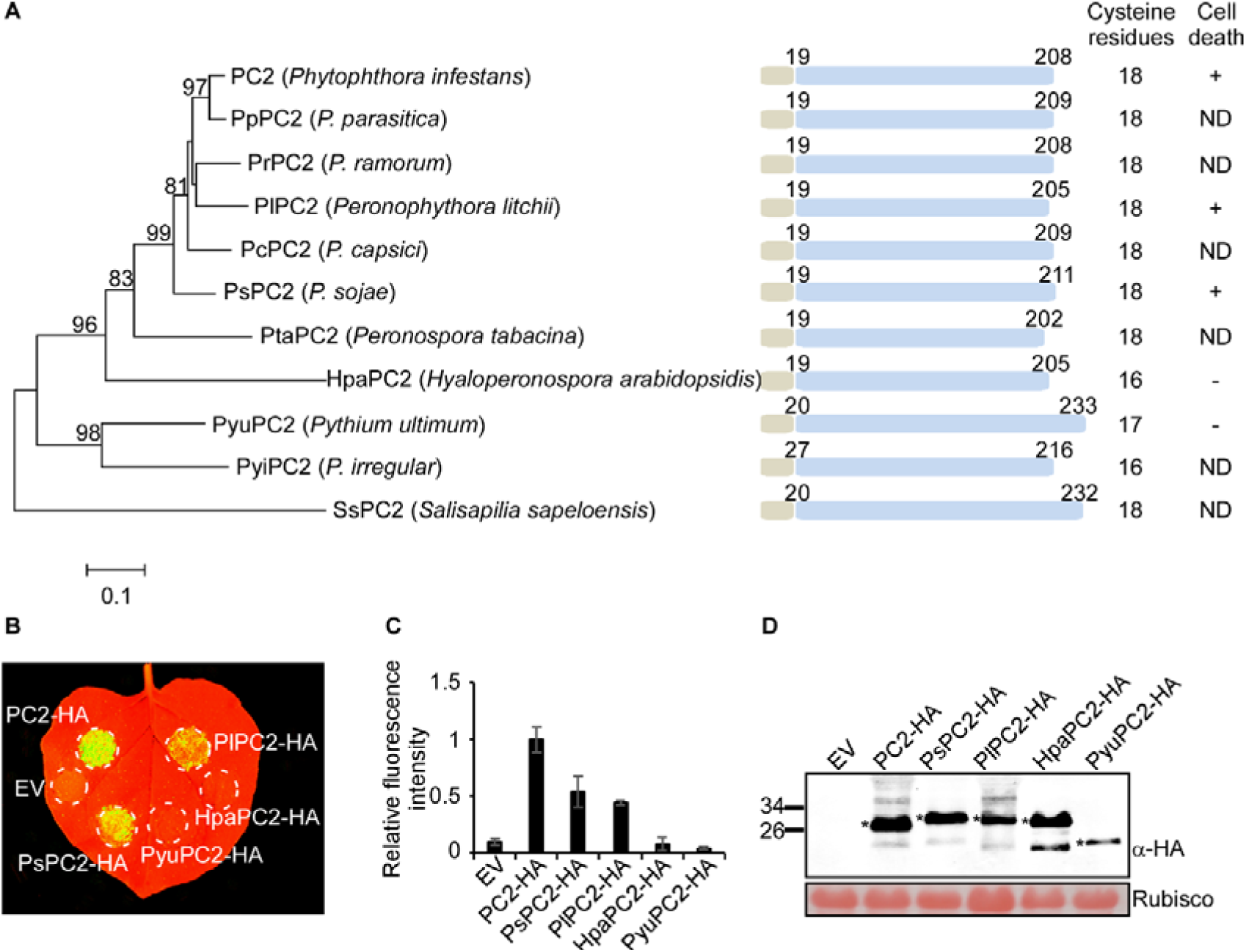
*PC2* homologs are widely present in oomycete species. (**A**) Phylogeny of PC2 from selected species. The tree was constructed by the neighbour-joining (NJ) algorithm implemented in MEGA 7. Bootstrap percentage support for each branch is indicated. The scale bar represents 10% weighted sequence divergence. “+”, cell death induction in *N. benthamiana*; “-”, no cell death induction in *N. benthamiana*; “ND”, not determined. (**B**) PC2 homologs from *Phytophthora sojae* (PsPC2-HA) and *Peronophythora litchii* (PlPC2-HA) induced cell death in *N. benthamiana*. (**C**) Quantification of the fluorescence intensity induced by PC2 homologs. The fluorescence intensity induced by PC2 was assigned the value 1.0. Means and standard errors from five independent replicates are shown. (**D**) Immunoblot analysis of PC2 or homologs protein fused with HA tags transiently expressed in *N. benthamiana* leaves. “*” indicates immuno-positive band with expected size.

To determine the function of the *PC2* orthologs, the orthologs from *P*. *sojae*, *Peronophythora litchii*, *Hyaloperonospora arabidopsidis* and *Pythium ultimum* were cloned and transiently expressed in *N. benthamiana*. The orthologs from *P*. *sojae* (PsPC2) and *P. litchii* (PlPC2) triggered visible cell death, while the orthologs from *H*. *arabidopsidis* (HpaPC2) and *P. ultimum* (PyuPC2) did not induce cell death symptoms (Figures 2B and 2C). Immuno-blot analysis using an HA antibody confirmed that PC2 and its orthologs were successfully expressed in *N. benthamiana* leaves (Figure 2D). This data indicates that the PC2 orthologs in oomycetes are conserved but show functional divergence.

### A core minimal peptide of PC2 (residues 138 to 198) and two conserved cysteine residues are required for PC2 activity

The conservation of sequence and the diversity in function of PC2 orthologs prompted us to identify the sequence signatures that are required for PC2-induced cell death. Based on the predicted secondary structure of the PC2 protein, a series of PC2 mutants designated from PC2^M1^ to PC2^M6^ were constructed and transiently expressed in *N. benthamiana* (Figure 3A). Through a fragment truncation assay, we narrowed down the functional region to between 138-198 amino acids (Figures 3B and 3C). The cognate PC2 mutant designated as PC2^M4^ elicited even stronger cell death when compared with the full-length *PC2*. In contrast, the PC2^M5^ or PC2^M6^ mutants, which were further reduced in size, lost full activity in triggering cell death (Figures 3B and 3C). Accordingly, we found the HpaPC2^M1^ mutant, the HpaPC2 corresponding peptide matching to PC2^M4^, does not have such function. To confirm the minimum PC2 functional peptide sequence, a HpaPC2 chimeric mutant HpaPC2^M3^ was created by substituting the HpaPC2 region (140-200) with the minimum functional fragment of PC2. This resulting PC2 mutant gained cell death activity (Figures 3B and 3C), suggesting that the 138-198 fragment of PC2 is pivotal to induce plant defense responses.

**Figure 3.**
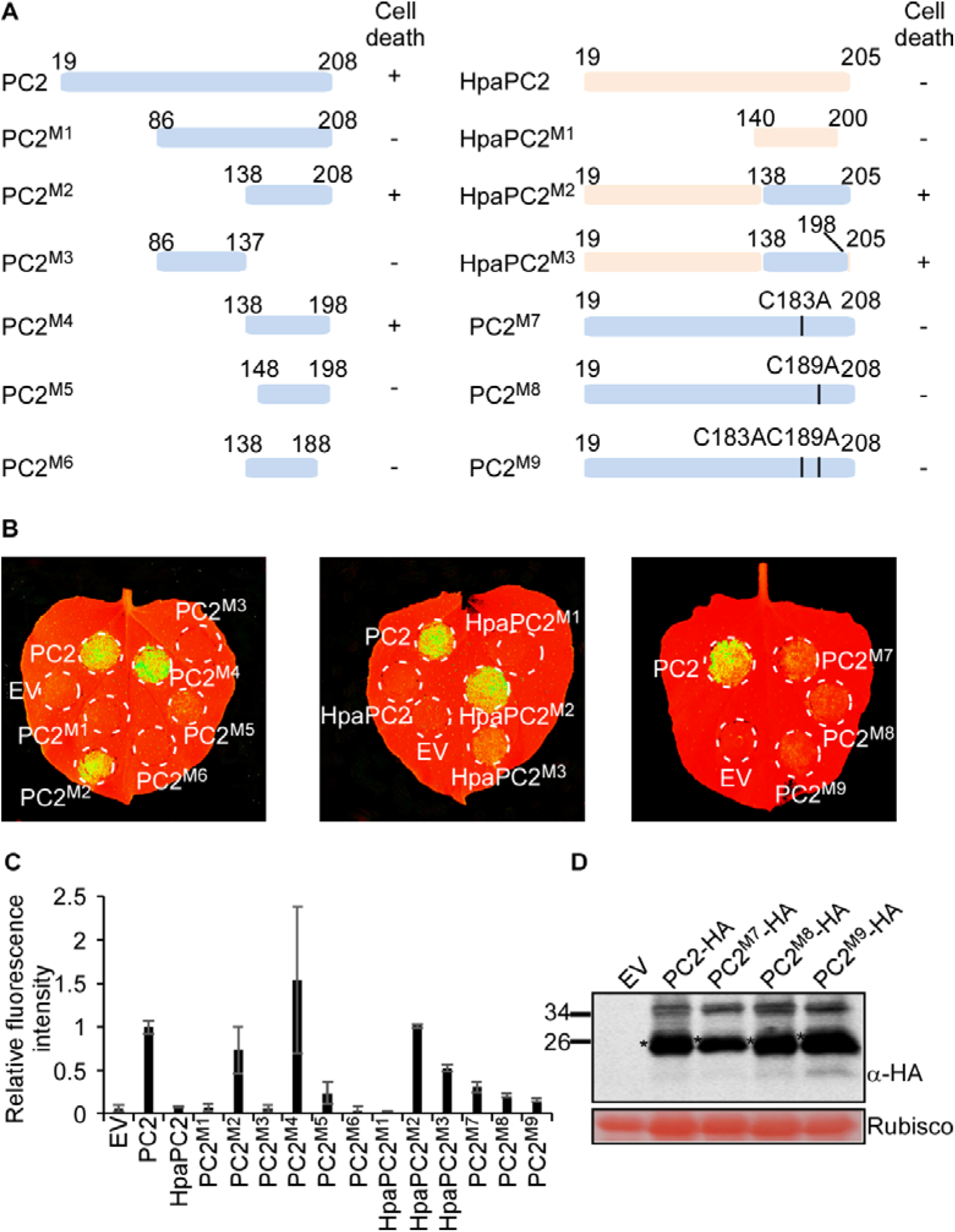
PC2 protein sequence signatures that are required for activity. (**A**) Identification of the functional peptide of PC2 for inducing cell death. Various PC2 mutants were constructed according to PC2 and HpaPC2 functional differentiation. “+”, induce cell death in *N. benthamiana*; “-”, cannot induce cell death in *N. benthamiana*. (**B**) Representative *N. benthamiana* leaves 5 d after inoculation with *Agrobacterium* strains carrying a series of mutants. (**C**) Quantification of the fluorescence intensity induced by different mutants in *N. benthamiana* leaves. The fluorescence intensity induced by PC2 was assigned the value 1.0. Means and standard errors from three independent replicates are shown. (**D**) Immunoblot analysis of PC2 mutants fused with HA tags transiently expressed in *N. benthamiana* leaves. “*” indicates immuno-positive band with expected size.

The PC2 minimal sequence contains seven cysteine residues (Supplementary Fig. 6A). We speculate that these cysteines play a role in maintaining effector structure and are essential for PC2-induced cell death. To test this hypothesis, we replaced these cysteines by a PCR-based site-directed mutagenesis alone or together with alanine in the PC2^M4^ minimal peptide and produced 8 mutants (PC2^M4(C146A)^, PC2^M4(C152A)^, PC2^M4(C161A)^, PC2^M4(C177A)^, PC2^M4(C183A)^, PC2^M4(C189A)^, PC2^M4(C198A)^, and PC2^M4(7CA)^). Cell death assays were carried out in *N. benthamiana* and substitution of the cysteine residue at position 183 or 189 was shown to impair cell death, suggesting these two residues are required for this function (Supplementary Figures 6B and 6C). To further confirm these results, we substituted cysteine residues at positions 183 and/or 189 with alanine in the full-length PC2 sequence and generated three mutants (PC2^M7^, PC2^M8^, and PC2^M9^). All mutants lost their cell death activity when tested in *N. benthamiana* (Figures 3B and 3C), although western blots confirmed the corresponding proteins remained stable (Figure 3D). Interestingly, the two cysteine residues at positions 183 and 189 are conserved in all PC2 orthologs. Thus, our PC2 mutagenesis assay demonstrates that the function of PC2 in triggering cell death in *N. benthamiana* requires a core minimal sequence from 138 to 198 and the two conserved cysteine residues.

### Plant serine protease P69B mediates cleavage of PC2 protein

We found in the previous experiment that the PC2^M1^ mutant does not trigger cell death whereas the shorter truncation mutants have functions (Figure 3B), so we hypothesized that PC2 protein requires further processing to generate bioactive peptides. To test this theory, we infiltrated a plant protease inhibitor cocktail into *N. benthamiana* following transient expression of PC2. This significantly impaired PC2-triggered cell death (Figure 4A). To figure out which component of the protease inhibitor cocktail caused the impairment, we infiltrated each individual components of cocktail into *N. benthamiana* leaves six hours before transient expression of PC2. Cell death was strongly inhibited by serine protease inhibitor 3, 4-Dichloroisocoumarin (DCI), whereas the other protease inhibitors such as cysteine protease inhibitor E-64 did not inhibit PC2-induced cell death (Figures 4A and 4B). Thereby, we reasoned that PC2 may be cleaved by serine proteases to produce bioactive peptides, which can be later recognized by the plants. To identify candidate plant proteases suppressed by DCI, we monitored the activity of serine proteases in the apoplast of tomato and *N. benthamiana* using the fluorophosphonate activity-based probe FP-TAMRA(Liu et al., 1999). Preincubation of the apoplast extract with DCI strongly suppressed labeling of 70 kDa subtilases in apoplastic fluid in both plant species (Figure 4C). This indicates that subtilases might be required for PC2-induced cell death, with P69B a likely candidate as it is the most abundant subtilase displayed with FP-TAMRA in apoplastic fluid of tomato(Sueldo et al., 2014).

**Figure 4.**
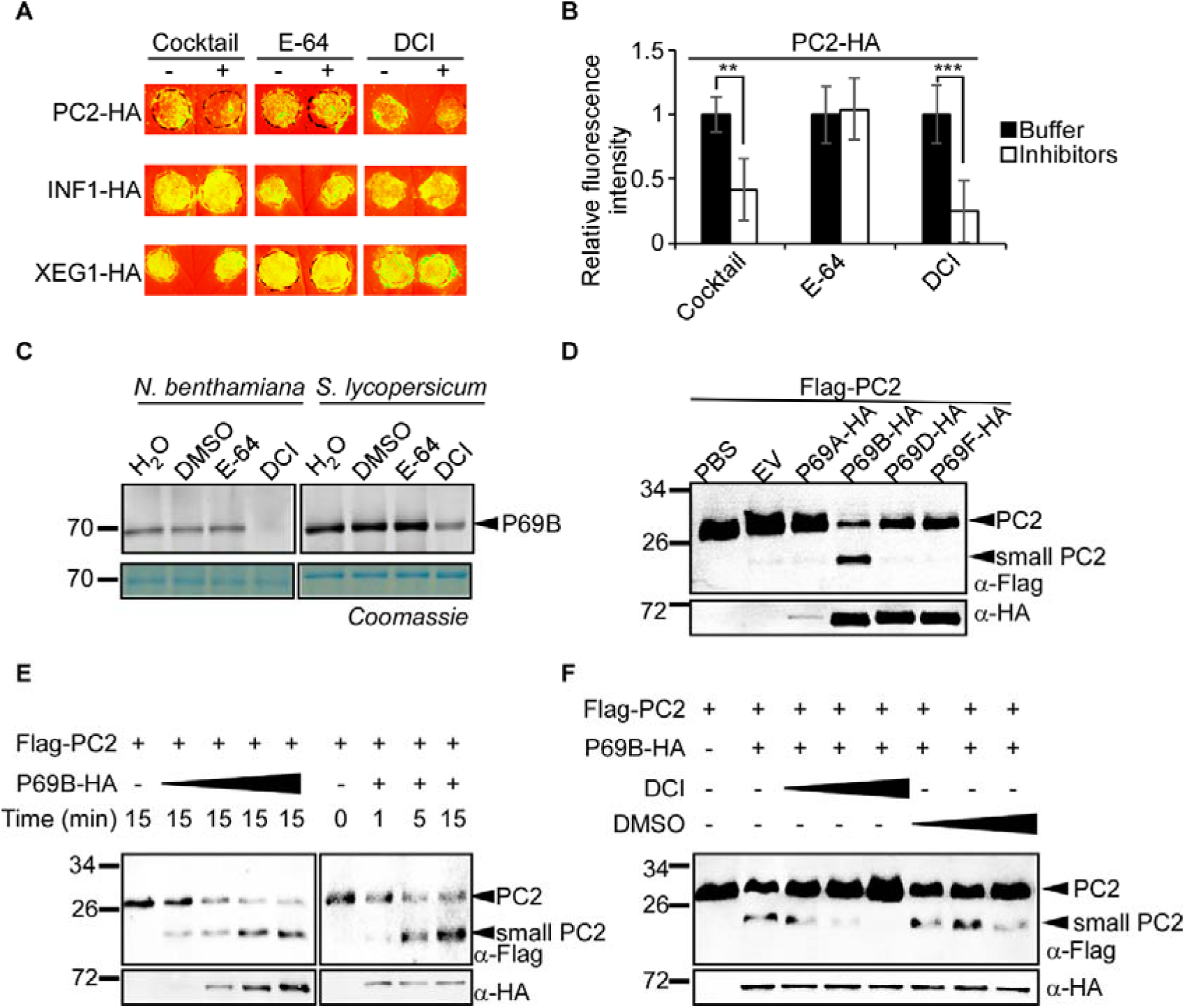
Plant subtilase P69B cleaves PC2 protein. (**A**) PC2-induced cell death in *N. benthamiana* leaves was inhibited by plant protease inhibitor cocktail and Serine protease inhibitor DCI. The *N. benthamiana* leaves were infiltrated with the cocktail and E-64 6 hours before agrobacterium-injected PC2, or DCI 12 hours after PC2 expression. E-64, Cysteine protease inhibitor, used as a control. (**B**) Quantification of the fluorescence intensity in *N. benthamiana* leaves after treatment by chemical inhibitors. Fluorescence intensity triggered by PC2 under respective buffer treatment was assigned the value 1.0. Means and standard errors are calculated from six replicates (**P< 0.01; one-way ANOVA). (**C**) DCI inhibits serine protease activity in tomato and *N. benthamiana* apoplastic fluids. The apoplastic fluids were incubated for 30 min with DCI. E-64 was used as a control. Active serine proteases were labelled using FP-Tamra. Labelled proteins were detected in a protein gel by fluorescent scanning. (**D**) The PC2 protein was cleaved by P69B in vitro. HA-tagged P69B or homologs were expressed in *N. benthamiana*, and FLAG-PC2-His protein was expressed in *E. coli*. PC2 and P69B or its homologs protein were co-incubated at 10°C for 30 minutes. The small band indicated PC2 cleavage products and were detected using anti-FLAG immunoblot. (**E**) Cleavage of PC2 was gradually increased after adding the various concentrations of P69B or the incubation time. (**F**) Serine protease inhibitor DCI prevented P69B cleavage of PC2 protein. HA-tagged P69B protein was added into the tube after FLAG-tagged PC2 protein and DCI were co-incubate in 4°C for 30 minutes. The cleavage products were gradually reduced in the presence of 0.5 mM, 1 mM, and 2 mM DCI. The cleavage products were detected using anti-FLAG immunoblot.

To explore whether the activity of P69B is required for PC2 function we performed an in vitro cleavage experiment. Recombinant PC2 protein with N-terminal FLAG tag (FLAG-PC2) was expressed and purified in *E. coli*. P69B and its homologs P69A, P69D, P69F fused to an HA tag were ectopically expressed in *N. benthamiana*. The apoplastic fluid from *N. benthamiana* leaves transformed with the empty vector control resulted in weak cleavage of PC2, but no such event was observed in PBS buffer (Figure 4D). This data indicates that PC2 can naturally be cleaved in *N. benthamiana* extracellular proteases. Apoplastic fluid fractions from P69B expressing leaves showed a very clear cleavage of PC2 that was not found when expressing P69A, P69D, or P69F (Figure 4D), suggesting that PC2 is specifically cleaved by P69B. Furthermore, we co-incubated recombinant PC2 with different concentrations of P69B extracted from *N. benthamiana* apoplastic fluid with different incubation times. The data showed that the degradation of PC2 was increased with increasing abundance of P69B or incubation time (Figure 4E). To confirm that P69B cleaves PC2 in a protease activity-dependent manner, we introduced the DCI inhibitor in a co-incubation *in vitro* experiment. We found that PC2 cleavage was attenuated with increasing DCI concentrations (Figure 4F). However, DCI solvent, DMSO only weakly affected PC2 cleavage at high concentrations. Together these data illustrate that the serine protease P69B cleaves PC2 *in vitro* in a protease activity-dependent manner.

### Serine protease P69B is required for PC2-triggered immunity

To address the question of whether PC2 cleaved by P69B is required for PC2-induced cell death activity, we tested PC2 activity on P69B silenced tomato plants(Paulus et al., In submission). Here, we used RT-qPCR to examine *P69B* expression and confirmed significant silencing of *P69B* in tomato line asP69B (Figure 5A). We then extracted the apoplastic fluids from asP69B and wild-type (WT) lines. Recombinant PC2 protein was cleaved when co-incubated with apoplastic fluid from WT plants but not when co-incubated with apoplastic fluid from *asP69B* plants (Figure 5B).

**Figure 5.**
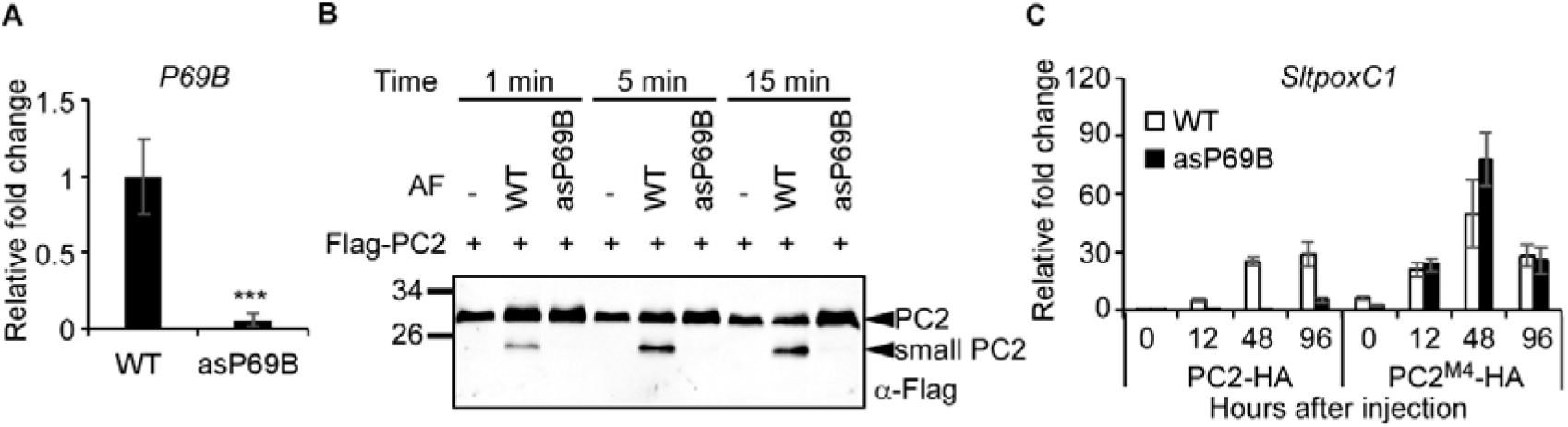
Plant subtilase P69B is required for PC2-triggered immunity. (**A**) The *P69B* expression level in asP69B tomato determined by qRT-PCR. Actin was used as an endogenous reference for normalization. Means and standard errors from three biological replicates are shown. Asterisks indicate significant differences (***P< 0.001; one-way ANOVA). (**B**) Apoplastic fluid extracted from asP69B did not cleave PC2 protein. PC2 recombinant protein was co-incubated with the apoplastic fluids (AF) extracted from WT or asP69B tomato plants. Cleavage products were detected using anti-FLAG immunoblot. (**C**) PC2-induced up-regulation of tomato PR gene *SltpoxC1* was impaired in asP69B plants. PC2^M4^-induced gene expression was not impaired in asP69B plants. The gene expression was examined at different time points by qRT-PCR.

To test if P69B is also required for PC2-induced cell death, we transiently expressed PC2 and the PC2^M4^ fragment in WT and asP69B tomato plants. PC2 induced cell death was very weak in WT tomato plants. We therefore performed RT-qPCR to detect the induction of a tomato defense marker gene *SltpoxC1*(Rivas et al., 2004) as an alternative method. We found that the tomato marker gene *SltpoxC1* was up-regulated in response to PC2 expression, however, this gene was not induced in the *asP69B* tomato line (Figure 5C). Interestingly, *SltpoxC1* was induced in both WT and asP69B tomatoes when these plants transiently expressed the PC2^M4^ mutant, providing further evidence that P69B participates in PC2 cleavage but not perception of PC2 in tomato. These data suggest that *P69B* silencing in tomato prevents cleavage of PC2 and consequently fails to generate a bioactive fragment to induce immunity.

Next we also detect whether the cleavage process is important for PC2 triggered immunity in *N. benthamiana*. We used VIGS to suppress the expression of a major apoplastic subtilase SBT5.2 in *N. benthamiana*(Paulus et al., In submission) and monitored PC2-induced cell death in the silenced plants. PC2-induced cell death was significantly weaker in *TRV:SBT5.2* plants when compared to TRV:*GFP* plants, but PC2^M4^ don’t show any difference (Supplementary Figures 7A and 7B). Immunoblotting confirmed that PC2 was successfully expressed at the expected size in *N. benthamiana* inoculated with *TRV:SBT5.2* or *TRV:GFP* (Supplementary Figure 7C). These data further confirm that the function of PC2 in triggering plant immunity is dependent on plant subtilases.

To further clarify that the cleavage process is important for PC2 triggered immunity, we generated a serious of PC2 mutants and tested their function and *in vitro* cleavage by P69B. Firstly, five PC2 mutants carrying large scale alanine substitutions were generated and only mutant PC2^103-116A^ was unable to induce cell death in *N. benthamiana* (Supplementary Figure 8A). Immunoblotting confirmed that PC2 mutants were successfully expressed in *N. benthamiana*, but over accumulation of PC2^103-116A^ is observed. In contrast, PC2 and another mutant PC2^116-126A^ that still trigger cell death maintain normal protein accumulation (Supplementary Figure 8B). Compared to PC2 and PC2^116-126A^, PC2^103-116A^ is resistant to cleavage by P69B in our *in vitro* cleavage assay (Supplementary Figure 8C). These results further confirm that the PC2 cleavage process is critical for its function and the cleavage site is very likely located in region 103-116.

### Serine protease inhibitors of *P. infestans* counteract PC2 cleavage

Earlier studies showed that *P. infestans* secretes Kazal-like protease inhibitor EPI1 and EPI1 homologs to repress P69B protease activity(Tian et al., 2004; Tian et al., 2005). We speculate that EPI inhibitors are able to suppress PC2-mediated plant immunity by directly preventing PC2 cleavage and PC2 recognition. To test this hypothesis, we selected six *P. infestans* genes encoding *EPI1*, *EPI2*, *EPI4*, *EPI6*, *EPI10*, and *EPI12* and cloned them into a plant expression vector. We found that transient expression of EPI1, EPI4 and EPI10 suppressed PC2-induced cell death but had no impact on INF1-and XEG1-induced cell death, suggesting PC2 inhibition by EPI inhibitors is specific (Figure 6A, Supplementary Figures 9 A-D). These data are consistent with our earlier observation that the chemical inhibitor DCI cannot inhibit INF1 and XEG1 mediated cell death (Figure 4A).

**Figure 6.**
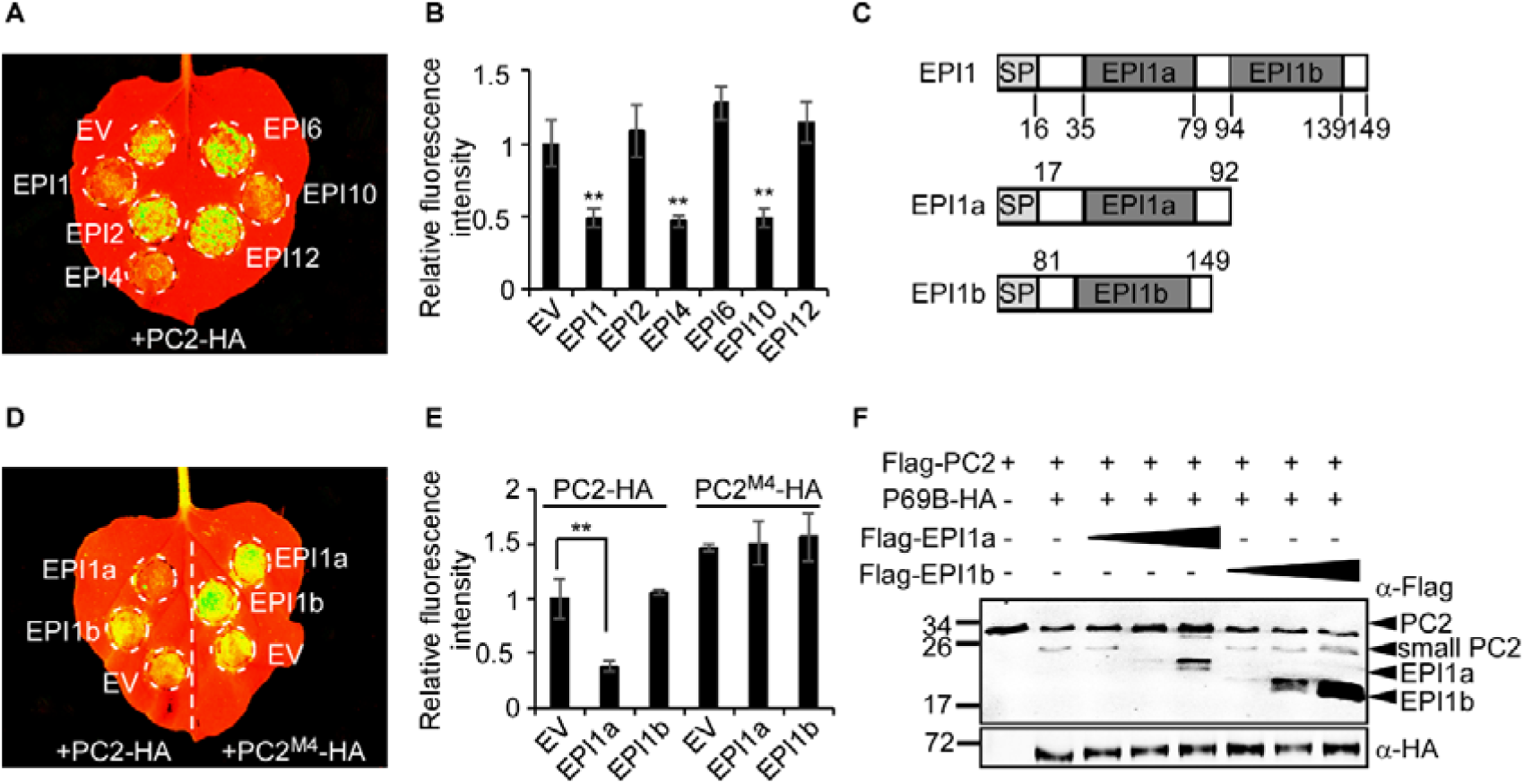
Serine protease inhibitors secreted by *P. infestans* counteract PC2 cleavage. (**A**) PC2 induced cell death in *N. benthamiana* leaves was inhibited by *P. infestans* secreted serine protease inhibitor EPI1, EPI4 and EPI10. The *N. benthamiana* leaves were infiltrated with different EPIs 36 hours before PC2 expression. (**B**) Quantification of fluorescence intensity induced by PC2 in *N. benthamiana* leaves under different EPIs treatment. The PC2 elicited fluorescence was normalized with empty vector (EV) treatment as reference. Means and standard errors were calculated from six replicates (**P< 0.01; one-way ANOVA). (**C**) Schematic representation of EPI1, EPI1a, and EPI1b that are used in the inhibition assay. The signal peptide (SP) and two Kazal domains (EPI1a and EPI1b) are shown in gray. EPI1a is a functional inhibition domain. (**D**) Cell death induced by PC2 was inhibited by EPI1a. The *N. benthamiana* leaves were infiltrated with the EPI1a and EPI1b 36 hours before PC2 expression. (**E**) Quantification of fluorescence intensity induced by PC2 in *N. benthamiana* leaves after EPI1a treatment. The PC2 elicited fluorescence was normalized with EV treatment as reference. Means and standard errors are calculated from six replicates (**P< 0.01; one-way ANOVA). (**F**) Recombinant EPI1a inhibits P69B cleavage of PC2 protein. FLAG-EPI1a and FLAG-EPI1b recombinant protein were expressed in *E. coli*. After the preincubation of P69B-HA with EPI1a or EPI1b, FLAG-PC2 protein was detected by immunoblot. FLAG-PC2 were gradually reduced in the presence of 0.5, 2.5, or 12.5 µM purified EPI1a protein. EPI1b, a non-functional Kazal-like domain, served as a control.

To examine whether EPI1 suppression of PC2 depends on its inhibitory function, we utilised mutant EPI1^P1^, an EPI1 mutant with the predicted active site P1 replaced with alanine(Tian et al., 2004) (Supplementary Figure 10A). Transient expression in *N. benthamiana* illustrated that EPI1^P1^ was unable to repress PC2-mediated cell death. However, both EPI1 and EPI1^P1^ did not inhibit PC2^M4^-induced cell death, indicating that the EPI1 inhibitor acts upstream of PC2^M4^ (Supplementary Figures 10B, 10C and 10D). EPI1 is composed of two Kazal-like domains, the four-cysteine atypical domain EPI1a, and the typical domain EPI1b(Tian and Kamoun, 2005). We cloned these two domains independently and tested their activity in repressing PC2-induced cell death (Figure 6C). PC2-triggered cell death was inhibited by EPI1a but not by EPI1b (Figures 6D and 6E). Neither EPI1a nor EPI1b repressed PC2^M4^ activity. This is in line with earlier reports indicating that EPI1a rather than EPI1b inhibits P69B protease activity.

We next examined whether the EPI1 protein could inhibit PC2 cleavage induced by P69B. Incubation of recombinant EPI1a with PC2 indicated that EPI1a did inhibit PC2 cleavage in a concentration-dependent manner and this inhibition was not observed with the non-functional EPI1b fragment (Figure 6F). These results confirm that subtilases play an important role in PC2 cleavage and that *P. infestans* protease inhibitors EPI1, EPI4 and EPI10 can block PC2 cleavage and PC2-induced cell death, depending on their inhibition activity.

## Discussion

More than 20 years ago, tomato subtilisin-like protease P69B was identified as a PR protein that is rapidly induced at early infection by citrus viroid(Tornero et al., 1996). P69B contains all the essential catalytic residues and is considered as an active protease that is involved in proteolytic defense in the plant extracellular matrix(Tornero et al., 1997). Tian and colleagues demonstrated that P69B is a direct target of EPI1, a Kazal-like serine protease inhibitor from *P. infestans*, suggesting P69B activity inhibition is important for pathogen colonization(Tian et al., 2004). However, the role of P69B in plant defense has remained elusive. Here we provide biochemical evidence that P69B proteolytically cleaves *Phytophthora* apoplastic effector PC2. *In vivo* inhibition of P69B through the application of a chemical inhibitor or EPI1 suppress PC2-triggered cell death (Figure 4A, Supplementary Figures 10B and 10C). In line with this observation, silencing of *P69B* in plants then prevents cleavage of PC2 and impairs PC2 activated PR gene up-regulation (Figures 5B and 5C). All these data indicate that PC2 proteolytic cleavage by P69B is critical for PC2 triggered immunity. This is likely because the cleavage of PC2 releases a short bioactive peptide which can be recognized by plant PRRs. This is supported by the fact that PC2-triggered immunity in *N. benthamiana* requires *BAK1*, an important co-receptor for LRR-RK-type PRRs.

Plant proteases present a broad spectrum of biological functions. In this study, we show that PC2 is targeted by the protease P69B. Tomato P69B paralogs such as P69A and P69D, contain all the catalytic residues but do not cleave PC2, suggesting the function of these family members is different. This could be due to gene duplication and subsequent neo-functionalization during evolution. To identify the substrates of P69B and paralogous proteases using biochemical approaches would help to fully understand the mode of action of these plant proteases. In general, plant proteases proteolytically cleave pathogen effectors to impair their biological functions and are considered important components of the defense response. Interestingly, a recent study of the *Ustilago* fungal protease inhibitor Pit2 demonstrated that pathogen utilizes plant extracellular protease to generate a powerful inhibitory peptide(Misas Villamil et al., 2019). In another study, *Arabidopsis* aspartic proteases cleave the evolutionarily conserved bacterial protein MucD to inhibit bacterial pathogen growth as an antibacterial strategy(Wang et al., 2019). Different from these discoveries, we showed here that extracellular cleavage of a *Phytophthora* effector PC2 by the plant protease P69B triggers immunity. This illustrates that plant extracellular proteases can be employed by both plants and their pathogens during co-evolution. Exploring the underlying mechanism is an important strp to designing rational plant disease management strategies.

Studies of plant proteases in other pathosystems have indicated that proteases bind and preferentially cleave their substrate in a sequence context favorite manner. For instance, it was recently shown that the P69A serine protease prefers to cleave residues after Asp(Reichardt et al., 2018). However, we predicted a few putative sites and the PC2 site-directed mutants we generated remained susceptible to P69B cleavage *in vitro*. Accordingly, these mutants still exhibited a cell death phenotype. We reason that either the P69B protease cleaves other sites or that the selection substrates may not be sequence dependent only. Other factors such as modifications or conformation of the substrate may impact the specificity between P69B and its substrates. Western blot data revealed that PC2 has a larger molecular weight than anticipated, suggesting modifications of PC2 *in vivo* may exist. Future work will aim to identify the cleavage sites on PC2 and to capture the bioactive peptides that act as signals to induce defense responses.

Ortholog searches revealed PC2 orthologs present in a wide selection of oomycete species including downy mildew and *Salisapilia species*, although no homolog was found in *Albugo* species. In particular, PC2 orthologs are highly conserved across *Phytophthora* species and all *Phytophthora* PC2 proteins tested were able to trigger cell death in *N. benthamiana*. In contrast, the downy mildew ortholog, HpaPC2, was highly divergent in sequence from *Phytophthora* PC2 and was unable to induce cell death in *N. benthamiana*. Whether they activate immunity in their respective host plants remains to be determined. Interestingly, 16 of the 18 cysteines present in *Phytophthora* PC2 are conserved in other oomycete species and substitution of just two of these cysteines (C183A and C189A) was sufficient to abolish PC2 activity. This suggests these two residues may form a potential disulfide bond to maintain protein conformation. However, both cysteines are conserved between functional and non-functional PC2 orthologs. We reason that downy mildew PC2 homologs failed to trigger immunity due to sequence variations beyond these two residues. Nevertheless, it appears that PC2 is conserved across the oomycetes and that plants may have evolved immune machinery to specifically recognize this pathogen signal. To evade this plant surveillance system, *Phytophthora* has evolved protease inhibitors such as the Kazal-like protease inhibitors (EPIs) to block PC2 extracellular processing. However, obligate biotrophic pathogens may evade recognition through loss of function mutations such as deletion of the *PC2* homolog in *Albugo* and gain of sequence polymorphisms as seen in downy mildew species.

Revealing the working mechanism of plant immunity to MAMPs provides solutions to enhance plant resistance. Whether PC2 plays a biological role during infection remains unknown. In this study, we were unable to knockout *PsPC2* in *P. sojae* by CRISPR/Cas9, suggesting PC2 may have a basic biological function that deserves further exploration. Overexpression of PC2 in *P. infestans* impairs infection on potato leaves, indicating that overexpression of PC2 may trigger an enhanced plant immune response against *P. infestans*. PC2 triggers cell death in several *Solanum* plants such as *N. benthamiana*, *N. tabacum*, *S. lycopersicum*, and *S. tuberosum*. Meanwhile, P69 homologs are present in an array of *Solanum* genomes. We therefore hypothesize that plant evolved PRRs recognizing PC2, and the recognition pathway may be conserved across *Solanaceae* species. This implies that the plant PRRs recognizing PC2 could be valuable in agricultural applications. Indeed, emerging studies reveal that genetic manipulation of plant PRRs recognizing pathogen extracellular effectors such as elicitin, NLP, XEG1, enhance basal resistance to *Phytophthora(Albert et al., 2015; Wang et al., 2018)*. Cloning of the specific *PRR* gene and functional allele screen may provide a resource to engineering durable plant resistance to *Phytophthora*.

## Materials and Methods

### Identification of candidate apoplastic effectors

The predicted secretome of *P. infestans* was previously defined(Raffaele et al., 2010a) and each of the 1,412 secreted proteins annotated herein based on features typical of known effector proteins. This included: (i) identification of Pfam domains using batch searches with default parameters(Sonnhammer et al., 1997), (ii) sequence similarity searches across the *Phytophthora* species cluster representing clade 1c species(Raffaele et al., 2010b), searches across *Phytophthora* species in general and searches against a list of core *P. infestans* orthologs(Haas et al., 2009) using BlastP searches with e-value cutoffs of 10^-5^, (iii) classification of small cycteine-rich proteins that were defined as those less than 200 amino acids in mature protein and having a cysteine number greater than or equal to 4 and a systeine content higher than 5%(Stergiopoulos and de Wit, 2009), (iv) sequence similarity searches across the predicted secretomes of an array of fungal plant pathogens (Supplementary Table 1) with e-value cutoffs of 10^-5^, (v) presence/absence of the RxLR motif using a custom perl script, (vi) analysis of gene expression during mycelium, sporangia, and infection stages(Raffaele et al., 2010a; Ah-Fong et al., 2017; Chen et al., 2018), (vii) sequence similarity analysis with species in the oomycete genus *Salisapilia* using BlastP searches with e-value cutoffs of 10^-5^, and (viii) presence of genes in gene sparse regions in the *P. infestans* genome as described previously(Saunders et al., 2012) (Supplementary Table 1). These classified as being expressed in hyphae and planta and encoding small cysteine-rich (SCR) proteins were selected for functional analysis as putative apoplastic effector candidates (Supplementary Table 2).

### Growth of plants and microbes

*Nicotiana benthamiana*, *N. tabacum*, *Solanum melongena*, *S. tuberosum* (Desiree), and *S. lycopersicum* lines Cf-0 (Moneymarker) were grown in pots containing the mixture of sterile soil and vermiculite at 25°C under a 16-hours-light / 8-hours-dark condition. *Escherichia coli* strains JM109 and DH5α were used to propagate all plasmids and DH5α was also used to express secreted proteins. *Agrobacterium tumefaciens* strains GV3101 was used for agroinfiltration in plants. *E. coli* and *A. tumefaciens* were cultured on LB medium with the corresponding antibiotic at 37°C and 30°C, respectively. *Phytophthora infestans* wild-type strain MX5-1 and transformed lines were routinely maintained in rye medium at 18°C under dark condition.

### Molecular cloning and bioinformatic analysis

Full-length *PC2* gene and its orthologs sequences with *PR1* signal peptide sequence amplified from cDNA of various species using high-fidelity PrimeSTAR HS DNA Polymerase (Takara), primers pairs and restriction enzymes (Supplementary Table 4) were cloned into the reconstructive PVX vector pICH31160 for immunity test in *N. benthamiana*. Serine protease *P69s* genes and biotic serine protease inhibitor *EPIs* were cloned into the pICH86988 vector. Tobacco Rattle Virus (TRV)-based vector pTRV1 and pTRV2e were used for gene silencing in *N. benthamiana*.

Plasmid pFLAG-ATS (Sigma) was used for protein expression in this study. The DNA sequence encoding PC2 mature protein were cloned into pFLAG-ATS vector for secreted protein in *E. coli*. The DNA sequence encoding protease inhibitor EPI1 and its P1 mutant were cloned into pFLAG-ATS to generate expression vector pFLAG-EPI1 and pFLAG-EPI1^P1^ as previously reported(Tian et al., 2004). The plasmid pFLAG-EPI1a and pFLAG-EPI1b for expressing the individual Kazal domains of EPI1 in *E. coli* were constructed following a previous publication(Tian and Kamoun, 2005). All the constructs we used in this study were listed in Supplemental Table 5.

Signal peptide prediction was performed using the SignalP 4.1 server (http://www.cbs.dtu.dk/services/SignalP/). Alignment of PC2 and its orthologs from other oomycete species was performed with BioEdit. Molecular phylogenetic analysis was performed using maximum likelihood method in MEGA (v7.0) (https://www.megasoftware.net/) with default parameters based on the Poisson correction model.

### RNA isolation and qRT-PCR

Total RNA was extracted from samples treated with different conditions using the PureLink™ RNA Mini Kit (Invitrogen), following the manufacturer’s protocol. Agarose gel electrophoresis was used for RNA integrity test. RNA yield was measured using the NanoDrop Micro Photometer (NanoDrop Technologies). The first-strand cDNA for qRT-PCR was synthesized from 100 ng total RNA using PrimeScript RT reagent Kit with gDNA Eraser (TaKaRa). For qRT-PCR, SYBR Green dye (Takara) was used according to manufacturer’s instructions. The qRT-PCR reactions were performed on an ABI Prism 7500 Fast real-time PCR System (Applied Biosystems, Foster City, CA, USA).

### *P. infestans* transformation and transformants detection

Full-length (including signal peptide) *PC2* cDNA was cloned into the *Phytophthora* transformation vector pTOR, driven by the Ham34 promoter(Judelson, 1991). Transformation of *P. infestans* was achieved using a modified PEG-CaCl2-lipofectin protocol(Judelson, 1991), modified as described in Avrova et al.(Avrova et al., 2008). Transformed *P. infestans* lines were maintained at 18°C in rye agar containing 25 µg mL^-1^ G418 antibiotic.

For PC2 protein expression in *P. infestans*, *PC2* transformants were cultured in amended lima bean (ALB) liquid medium. The wild type strain was used as a negative control. Mycelium was harvested by centrifugation at six dpi, and total RNA was extracted from mycelium. Transcript levels were measured by qRT-PCR using the *P. infestans* beta-tubulin gene as an internal reference.

### Plant inoculations and relative biomass assay

The virulence assay of *P. infestans* transformants was carried out using *N. benthamiana* and susceptible potato leaves. The wild type and transformants were grown on rye agar without G418 selection for seven days before inoculation. Lesion size was measured at five days post inoculation. Primers specific for *P. infestans* beta-tubulin gene and *N. benthamiana* actin gene were used to quantify the relative biomass of *P. infestans* by qRT-PCR.

### Transient expression in *N. benthamiana*

*Agrobacterium tumefaciens*-mediated transient expression was performed using 4-week-old *N. benthamiana* plants. To test whether PC2 and its orthologs trigger cell death in *N. benthamiana*, *A. tumefaciens* strains GV3101 carrying PC2 or its orthologs cloned in pICH31160 vector and the P19 silencing suppressor, were infiltrated into *N. benthamiana* leaves. The number of necrotic leaves was scored, and the pictures were taken at 5 d post agroinfiltration.

VIGS experiments were conducted using *A. tumefaciens* strains harboring the pTRV1 and the corresponding pTRV2 vector. The agroinfiltration was performed on two-week old *N. benthamiana* plants, and the silenced plants (four-week old) were used for the cell death test. The silencing efficiency of target genes was measured using qRT-PCR with their expression values normalized to the level in TRV:*GFP* and using actin as the internal reference.

### Cell death analysis

*N. benthamiana* leaves were exposed under UV light (365 nm) in ultraviolet imager (Tanon 5200 Multi) at five days post agroinfiltration. The raw image was used for cell death intensity statistics using ImageJ software and for coloring by TanonImage software. Each treatment had at least three replications.

### Oxygen burst detection

Oxygen burst was observed based on H_2_O_2_ accumulation after staining *N. benthamiana* leaves with diaminobenzidine (DAB). Leaves at two days post agroinfiltration were collected into a solution containing 1mg mL^-1^. DAB dissolved in deionized water to which HCl was added to bring the pH to 3.8 to solubilize DAB. Leaves were then incubated in the growth chamber for an additional 8 h period to allow DAB uptake and reaction with H_2_O_2_ and peroxidase. The images were captured using a Canon digital camera at the designated time points and the DAB staining intensity on photographs was quantified using ImageJ.

### Extraction of the recombinant protein from plant tissues

*N. benthamiana* leaves expressing the recombinant proteins were ground to powder in liquid nitrogen. The powder was added to an extraction buffer (10% glycerol, 25 mM Tris pH 7.5, 1 mM EDTA, 150 mM NaCl), 10mM dithiothreitol (DTT), protease inhibitor cocktail (1:100, Sigma) and 0.15% NP-40 (Beyotime Biotechnology). Samples were centrifuged at max speed for 10 min at 4°C, and the supernatant was used for SDS-PAGE gel electrophoresis. Apoplastic fluids extracted from *N. benthamiana* leaves were prepared according to the method of de Wit and Spikman(Krüger et al., 2002). PBS (pH=5.5) was used as the solvent of protein P69B. For tomato leaves, a 0.24 M sorbitol solution was used as an extraction buffer(de Wit and Spikman, 1982). The apoplastic fluids were harvested for SDS-PAGE and other activity assays.

### Expression and enrichment of recombinant proteins from *E. coli*

*E. coli* strain DH5α containing the trans-genes cloned in pFLAG-ATS vector was used for protein expression(Kamoun et al., 1997). *E. coli* cultures were scaled up in different volume depending on the requirement. Culture filtrate was harvested based on the protocol described in Dong et al(Dong et al., 2014). The recombinant proteins with either FLAG-tag or His-tag were purified by corresponding immune affinity agarose beads (Sigma).

### SDA-PAGE and immunoblotting

Proteins were separated on 12% or 15% SDS-PAGE gels run at 80-120 V for 2 hr and electroblotted on PVDF membrane (Millipore) at 85 V for 3 hours at 4 °C. Electroblotted membranes were rinsed in PBST (PBS with 0.1% Tween 20) containing 5% (w/v) nonfat milk powder for 30 to 60 min. Primary antibodies were diluted in PBST solution to the ratio of 1:5000 and incubated with membranes for 3 hours at room temperature. Membranes were washed 3 times in PBST for 5 minutes in PBST before incubation with secondary antibodies (1:10000 dilution) for 0.5 h. After washing for 5 min thrice in PBST, signals were visualized by excitation at 800 nm following the manufacturer’s instructions (Odyssey, no. 926-32210; Li-Cor). The proteins expressed in *N. benthamiana* or *E. coli* were assessed by immunoblotting with anti-HA or anti-FLAG primary monoclonal antibody (Abmart).

### *In vitro* PC2 proteolytic cleavage assay

The in vitro cleavage test of PC2 was performed as follows. In planta-expressed recombinant P69B protein was extracted in PBS (pH5.5) using vacuum infiltration/centrifugation technique, and 10 µl extract was incubated with recombinant PC2 (1 µM) purified from *E. coli* in 50 µl volume at 10°C for 30 min. The reaction products were detected by western blot using anti-FLAG and anti-HA antibody.

### *In vitro* and *in vivo* protease inhibition assay

We tested the effect of several inhibitors to PC2-triggered cell death in *N. benthamiana* leaves. Each chemical inhibitor was infiltrated into *N. benthamiana* leaves 6 hours before PC2 agroinfiltration. Inhibitors used in this work included DPI (Sigma, 10 µM) dissolved in 0.1% DMSO, Protease inhibitor cocktail (Sigma, 1%) dissolved in 1% DMSO, and 3,4-Dichloroisocoumarin (DCI, Sigma, 1 mM) dissolved in 0.5% DMSO. For other inhibitors, Lacl_3_ (Sigma, 10 mM), AEBSF (Sigma, 1 mM), E-64 (Sigma, 10 µM) were dissolved in H_2_O. For inhibition assay of EPI effectors (EPI1, EPI2, EPI4, EPI6, EPI10, EPI12) from *P. infestans*, we infiltrated *Agrobacterium* strains carrying PC2 36 hours after EPIs agroinfiltration.

Inhibition of serine protease activity in tomato and *N. benthamiana* apoplastic fluids were assessed using a probe for serine proteases. First, 50 µL apoplastic fluid was pre-incubated with DCI (1 mM), E-64 (10 µM) or the same volume of DMSO for 30 min. Then, labelling of active serine proteases was performed by incubating 43 μL apoplastic fluid with 5mM sodium acetate (NaOAc), 5 mM DTT and 0.2 μM FP-Tamra (Thermo) for 1 h at room temperature in the dark. After incubation, the labelled proteins were acetone-precipitated and separated by 14% SDS-PAGE. Labelled proteins were visualized by in-gel fluorescence scanning using a Typhoon FLA 9000 scanner.

For *in vitro* proteolytic cleavage assay of PC2 by P69B, a broad-spectrum chemical serine protease inhibitor DCI was used to block P69B activity. The DCI was pre-incubation with PC2 protein 30 min before P69B protein was added into the tubes. To detect the inhibitor activity of EPI1a *in vitro*, we incubated purified EPI1a or EPI1b with P69B for 30 min followed by incubation with PC2 for 30 min at 10°C. The reaction products were detected by immunoblotting with anti-FLAG and anti-HA antibodies as described above.

## Author Contributions

S.W., R.X., S.F., J.P., M.S., J.W. and S.D. performed experiments; H.S., Y.W., V.V. and D.S. analyzed data; S.W., Y.W., X.Z., R.H., S.K. and S.D. designed experiments; S.W., Y.W. and S.D. wrote the manuscript; all authors commented on the article before submission.

## Acknowledgments

We are grateful for helps from Sylvain Raffaele, Abbas Maqbool, Jiorgos Kourelis, Chih-Hang Wu and Liliana M Cano in Kamoun’s group at The Sainsbury Laboratory and all the lab members in Dong’s group in NAU. Prof. Wenbo Ma (UCR) and Prof. Gunther Doehlemann (Univ. Cologne) are appreciated for helpful suggestions. We also thank Prof. Juyou Wu (NJAU) and Prof. Chuanyou Li (CAS) for offering plant materials. This work is supported by the National Natural Science Foundation of China (31772144, 31721004), the Fok Ying-Tong Education Foundation (151025), ERC Consolidator Grant 616449 and BBSRC grant BB/S003193/1. S. Wang received support from the Postgraduate Research & Practice Innovation Program of Jiangsu Province (KYCX17_0579).

